# Identification of first-in-class small molecule potential inhibitor of GDF15

**DOI:** 10.64898/2025.12.12.693902

**Authors:** Antonio Chiariello, Lorenzo Lenti, Lorenzo Trofarello, Manuela Sollazzo, Mattia Lauriola, Alberto del Rio, Stefano Salvioli, Marco Daniele Parenti, Maria Conte

## Abstract

**Background:** Growth Differentiation Factor 15 (GDF15), a mitokine implicated in stress response, has been associated with numerous diseases, particularly cancer cachexia. Many approaches, including monoclonal antibodies and peptide antagonists, have been implemented to inhibit GDF15 activity. It is currently unknown whether it is possible to inhibit GDF15 using small organic molecules (SOMs).

**Methods:** A structure-based *in silico* screening workflow of a curated compound library was implemented to identify SOMs capable of binding to the monomeric or dimeric form of GDF15. The three top-ranking SOMs of each group were then tested *in vitro* on normal and cancer cells.

**Results:** Among all tested SOMs, dioxoimidazolidin derivative named SOM D resulted capable of inhibiting GDF15 dimer formation and decreasing binding to GFRAL. Furthermore, it was found to be devoid of acute cytotoxicity both in normal cells (dermal fibroblasts, DFs) and tumor cells (OV90), to significantly slow proliferation and modify the expression of genes involved in GDF15 signaling or in cell cycle and senescence in OV90, but not in DFs. Interestingly, simultaneous treatment with SOMs and doxorubicin (doxo) failed in arresting the cell cycle compared to doxo alone.

**Conclusions:** The *in silico* screening has proven effective in identifying SOMs putatively capable of inhibiting GDF15, among which SOM D appears the most promising. Further characterizations will be necessary to better understand the exact mechanisms of action. Furthermore, the observation that SOMs have an opposite effect on proliferation in cancer versus normal cellular context suggests a dual role in cell-cycle control in presence of cancer aberration.

## Introduction

Growth differentiation factor 15 (GDF15), initially described as a distant member of the TGF-β superfamily, was discovered by three different research groups in 1997 (**Bootcov et al, 1997; Hromas et al, 1997; Lawton et al, 1997).** GDF15 is a stress-response molecule, expressed in different tissues, in particular in response to mitochondrial stress, and for this reason is considered a “mitokine”, a molecule secreted in tissues and cells with mitochondrial dysfunction, that can have beneficial or, possibly, detrimental effects also in tissues far from the site of production (**Durieux et al, 2011; Breit et al, 2021**). In the last years, GDF15 has also emerged as a key mediator in aging and age-related diseases. In fact, several studies found that GDF15 is one of the most upregulated circulating proteins during aging (**Tanaka et al, 2018; Conte et al, 2019; Lehallier et al, 2019).**

GDF15 gene is located on chromosome 19 and encodes for a 308 amino acid protein. The protein is first synthetized as a precursor (pro-GDF15) that dimerizes and is then cleaved at a conserved furine-like cleavage site (RXXR) to form the C-terminal mature GDF15 dimer (m-GDF15). This dimer is the form most commonly found in the bloodstream (**Conte et al, 2022; Wan and Fu, 2024).** The precise mechanism of GDF15 action is not completely understood. The only certain specific receptor identified for GDF15 is the GDNF α-like receptor (GFRAL), discovered in 2017 by four different research groups. GFRAL requires the action of the co-receptor REarranged during Transfection (RET). While RET is widely expressed across different tissues, GFRAL is believed to be mainly expressed in the nucleus of the solitary tract (NST) and the area postrema (AP), two small areas in the hindbrain (**Emmerson et al, 2017; Hsu et al, 2017; Mullican et al, 2017; Yang et al, 2017**); lower concentrations of this receptor have been observed also in other tissues like testis and adipose tissue as well as in some cancers (**Wang et al., 2019).** The interaction of GDF15 with GFRAL and RET activates different signaling pathways such as ERK1/2, Akt, FOS, and PLC-γ (**Takahashi, 2022).** The main biological activities of the GDF15-GFRAL axis include the regulation of body weight and the reduction of food intake (**Hüllwegen et al, 2025).** Moreover, GDF15 can also act on peripheral tissues by increasing lipolysis and oxidative metabolism (**Breit et al, 2021).** However, several actions of GDF15 not dependent on the activation of the neurons of the NST and AP have been reported. In particular, different studies proposed a role of GDF15 in regulating inflammatory processes. For instance, it has been shown that GDF15 could reduce the production of pro-inflammatory cytokines in liver of mice with fibrosis. Moreover, GDF15 KO in mice caused the overexpression of TNF-α by T cells and the activation of these cells, worsening liver injury (**Chung et al, 2017).** Another study showed that GDF15 was overexpressed in patients with sepsis and its activity was pivotal for survival bacterial and viral infections in mice (**Luan et al, 2019)**. On the other side, different studies showed a possible pro-inflammatory role for GDF15. In particular, a study showed an involvement of this mitokine in the progression of atherosclerosis, through an IL6-dependent inflammatory mechanism (**Bonaterra et al, 2012)**. GDF15 is also considered a member of the senescence-associated secretory phenotype (SASP) (**Di Micco et al, 2021)**. Furthermore, a role of GDF15 in cell cycle regulation has been proposed. It has been observed that GDF15 treatment increased cell proliferation of human umbilical vein endothelial cells (HUVECs) by enhancing the expression of G1 cyclins, cyclins D1 and E, through the PI3K/Akt, ERK, and JNK pathways (**Jin et al, 2012**).

GDF15 has been associated with several pathologies. In particular, an association has been found between GDF15 and type 2 diabetes (T2D), sarcopenia, cardiovascular diseases, hypertension, neurodegenerative diseases and renal dysfunctions **(Conte et al, 2022)**. For instance, we found a higher protein expression of m-GDF15 in the brain of Alzheimer’s patients as well as higher plasma level in patients with muscle atrophy, compared to controls **(Chiariello et al, 2023; Chiariello et al, 2024).** Moreover, an association of GDF15 levels with frailty and overall survival has been observed (**Conte et al, 2019; Conte et al, 2022; Conte et al 2025)**. GDF15 has also been associated with many types of cancer and cancer cachexia, in particular it has been associated with increased cancer migration and invasiveness, although results are conflicting and the role of GDF15 in cancer appears to be context-dependent **(Siddiqui et al, 2021).** However, GDF15 plasma levels are often found higher in cancer patients, suggesting that it could be considered both a biomarker and a potential target for treating different types of cancer **(Hüllwegen et al, 2025**). Finally, the role of GDF15 in inducing cancer cachexia is well established. Different studies in mice have shown that GDF15 mediates cancer cachexia, leading to weight loss, lean and fat mass loss, reduced food intake and cachexia, through the GDF15-GFRAL axis **(Hüllwegen et al, 2025).** A recent study showed that GDF15 determined weight loss by increasing energy expenditure in muscles (**Wang et al, 2023).** All the findings on GDF15-cancer relations increased the interest in developing inhibitors of GDF15 or GFRAL. In particular, pharmacological inhibition the GDF15-GFRAL axis could be useful to improve cancer cachexia and anorexia symptoms. For example, a phase 2 clinical trial for ponsegromab, a GDF15-blocking antibody, showed that patients with cancer cachexia had increased weight gain as well as improved appetite and physical activity following 12 weeks of treatment **(Groarke et al, 2024)**. Moreover, another trial showed that the addition of GDF15-blocking antibody to immunotherapy could help alleviate resistance to the immunotherapy (**Melero et al, 2025).** However, monoclonal antibodies have limitations, such as high costs, poor tissue penetration, safety concerns and less modularity of the effect. Recent studies have also reported peptide-based inhibitors targeting the GDF15/GFRAL axis. One investigation described C-terminal fragments of GDF15 which bound the GFRAL extracellular domain and inhibited receptor signaling in the micromolar range **(Alexopoulou et al., 2023)**. Another work developed a 29-residue peptide antagonist (“GRASP”) of the GFRAL–RET complex, that bound GFRAL and attenuated GDF15- and cisplatin-induced anorexia *in vivo* **(Borner et al., 2023)**. More recently, an article described bicyclic tandem peptides derived by phage-display and structure-guided design to mimic the GDF15 homodimer and inhibit signaling via the GDF15–GFRAL–RET complex **(Noisier et al., 2025)**. Although these peptidic inhibitors provide important proof-of-concept for antagonizing the GDF15–GFRAL interaction, a small-molecule inhibitor may offer several advantages, such as improved oral bioavailability, enhanced tissue penetration (including potential access to the central nervous system where GFRAL is expressed), greater metabolic stability, simpler manufacturing and formulation, and the ability to cross cell membranes or modulate allosteric sites rather than simply competing at the large surface protein-protein interface of the ligand–receptor. Thus, a well-designed small-molecule binder of GDF15 (or GDF15–GFRAL interface) could provide a more drug-like profile for therapeutic development. In this study, we performed an *in silico* screening on commercially available compounds in order to identify small organic molecules (SOMs) putatively able to bind and inhibit GDF15. We have analyzed the *in vitro* effects of six of such SOMs, in both normal and cancer human cells, in order to verify whether any of them were actually capable of inhibiting the action of GDF15. Results suggest that the dioxoimidazolidin derivative named SOM D seems to be the more promising for further testing.

## Materials and Methods

### Protein Structure Preparation

The crystallographic structure of human Growth Differentiation Factor 15 (GDF15) was retrieved from the RCSB Protein Data Bank (PDB Code 5VT2). Two biologically relevant oligomeric states were considered: monomeric and dimeric GDF15. Since no experimental evidences were available about possible active sites of GDF15 in both single-chain and dimeric forms, to define the most plausible binding regions distinct approaches were employed for the monomeric and dimeric protein forms. For the monomeric GDF15, the putative active site was selected based on intramolecular recognition features inferred from the protein’s own secondary structure.

Specifically, a short peptide fragment corresponding to residues 57–60 of the GDF15 sequence was used as a structural reference to locate the interaction region between the two monomers and therefore used to define the docking grid box for subsequent virtual screening. For the dimeric GDF15, the putative active site was identified using P2Rank server (**Polak et al., 2025**), a machine learning-based pocket prediction algorithm that evaluates geometric and physicochemical properties of the protein surface. The highest-ranked predicted pocket, located at the interface region between the two monomeric subunits, was selected for docking grid definition.

All protein structures were prepared using protein preparation workflow as included in the software package Maestro (Schrödinger Release 2024-2: Protein Preparation Workflow; Epik, Schrödinger, LLC, New York, NY, 2024; Impact, Schrödinger, LLC, New York, NY; Prime, Schrödinger, LLC, New York, NY, 2025). The preparation included removal of all crystallographic water molecules and heteroatoms not involved in ligand binding, correction of incomplete side chains, and addition of polar hydrogen atoms, as well as assignment of correct protonation states at pH 7.4.

### Ligand Library Preparation

The Hit Locator Library from Enamine (https://enamine.net/compound-libraries/diversity-libraries/hit-locator-library-460) was used for screening. Structures were downloaded in SDF format and converted to 3D conformers using Open Babel. Each ligand was subjected to energy minimization with the MMFF94 force field and assigned Gasteiger partial charges. To ensure a uniform and relevant screening set, duplicates, reactive compounds, and unstable species were filtered out using RDKit. The final ligand library contained approximately 460K unique structures, all stored in PDBQT format compatible with AutoDock Vina.

### Docking Protocol

Molecular docking was performed using AutoDock Vina v1.2.7 (**Trott & Olson, 2010; Eberhardt et al., 2021**) on both monomeric and dimeric GDF15 models. Docking grids were centered on the putative binding pocket identified as described above. The grid dimensions were adjusted to fully encompass the binding site, with spacing set to 1.0 Å. For each ligand, up to 20 poses were generated, and the best-scoring conformations were selected according to the Vina scoring function (an empirical estimate of binding free energy). Docking simulations were executed on a high-performance Linux environment using multicore parallelization to increase throughput.

### Post-Docking Analysis

For the 50 top-ranked compounds obtained from the docking, the residues predicted to interact with each ligand (based on the best-scoring binding pose) were extracted and tabulated in a binary interaction matrix. In this matrix, columns correspond to individual GDF15 dimer residues, while rows represent ligands, with each cell indicating the presence or absence of an interaction between a residue and a given compound. To evaluate the diversity of binding profiles, pairwise similarity between ligands was calculated using an asymmetric similarity metric defined as the proportion of residues with positive interaction values in the reference ligand that were also contacted by the compared ligand. This measure, computed across all 18 residues, quantifies the degree to which the interaction pattern of one compound is encompassed by another. By applying this criterion, the compounds displaying the most distinct interaction profiles were identified, ensuring maximal structural and functional diversity among the selected candidates.

### Compound Source

Selected molecules were purchased in milligram quantities from Enamine. Purity of compounds was ≥95%, as declared by the chemical vendor.

### Cell cultures, *Gdf15 in vitro* knock down (KD) and treatments

A cancer cell line and a normal cell line were used as *in vitro* models. As far as normal cells, commercially available primary dermal fibroblasts (DFs) (Cti biotech) obtained from 2 young donors (age range 24-30 years) were used. As far as cancer cells, OV90, a cell line derived from ovary papillary serous adenocarcinoma, were used. Both DFs and OV90 were cultured in high glucose DMEM supplemented with 10% heat-inactivated fetal calf serum (FCS), penicillin (100 units/ml), streptomycin (100 mg/ml), and 2 mM L-glutamine (all from Sigma), in an incubator at 5% CO_2_, with a humidified atmosphere of 37°C. The DFs used for the experiments were between the 8th and 12th passages.

GDF15 KD on OV90 was obtained using RNA interference (RNAi) strategy. Small interfering RNA (siRNA) targeting GDF15 and scramble siRNA (negative control) were purchased from Cohesion Biosciences. A specific combination of siRNA against GDF15 was selected after testing the silencing efficacy of different combinations of three different siRNA. Transfections were performed using ScreenFect siRNA reagent (ScreenFect GmbH), following manufacturer’s indications. Briefly, 70,000 cells were seeded in a 12-wells plate and scramble or GDF15 siRNA were mixed to transfection reagent, incubated at room temperature for 20 minutes and then added to the cells. Medium was replaced after 24h with fresh complete medium. Cells were harvested after a further 72h for RNA extraction.

The SOMs were resuspended in DMSO to make a stock solution of 50mM. A working solution of 1mM was obtained from the stock by diluting it in fresh complete DMEM. Treatments were performed in 24-well plates by seeding 35,000 cells/well. SOMs were added to the cells the day after seeding. Co-treatment with doxorubicin (NeoBiotech) was conducted incubating OV90 cells with 1µM of doxorubicin for 24h. After this 24h-period, the medium was replaced with fresh complete medium and 100μM of the SOMs were added. At the end of the treatment (24, 48 or 72h) cells were harvested, and viability and proliferation were evaluated with the Cell Drop FL cell counter (DeNovix) after staining with Trypan blue dye. Cells were then frozen at −80°C until RNA or protein extraction.

### RNA extraction and Real Time RT-PCR analysis

Total RNA was extracted from DFs and OV90 pellets using the EasyPure RNA kit (TransGen Biotech Co.). RNA quantification and purity analysis were performed using NanoDrop One Spectrophotometer (Thermo Scientific). cDNA synthesis was performed with HIScript III RT SuperMix for qPCR (+gDNA wiper) (Vazyme Biotech).

Gene expression was analyzed by Real Time RT-PCR, performed using SsoAdvanced Universal SYBR Green Supermix (Bio-Rad) and a Rotor gene Q 6000 system (Qiagen). A relative quantification was obtained using *Gapdh* as reference gene. The relative expression ratio was obtained using the 2^−ΔΔCT^ method. All the primers used in this study were predesigned and pre-validated and purchased from Bio-Rad. More information about primers is available at www.bio-rad.com/PrimePCR.

### Protein extraction and western blotting analysis

Protein extraction from OV90 was performed using RIPA buffer with the following composition: Tris HCl pH 8 50 mM, NaCl 150 mM, Sodium deoxycholate 0.5%, SDS 0.1% and Triton X-100 1%. Protease and phosphatase inhibitors (Sigma) were added to the lysis buffer. Pellets were resuspended in the buffer by vigorous pipetting, left on ice for 15min, vortexing few times. Lysates were then centrifuged at 15,000 rpm at 4°C for 15min and the supernatants were collected. Bradford’s method was performed for quantification, and purified proteins were then stored at - 80°C until used.

Protein expression was then analyzed by western blotting. 20µg of proteins were separated on a 4–15% Mini-PROTEAN^®^ TGX™ Precast Protein Gels (Bio-Rad), transferred on a PVDF membrane (Trans-Blot transfer, Bio-Rad) and then immunoblotted with the adequate primary antibody. GAPDH was used as loading control. Chemidoc system (Bio-Rad) was used for acquisition. Densitometric analyses of western blotting bands were performed using the Fiji software. Primary antibodies and dilutions used are listed in **Table 1**.

**Table 1.** List of primary antibodies used.

| Primary antibody and catalog number | Dilution used | Supplier |
| --- | --- | --- |
| GDF15 (ab206414) | 1:1000 | Abcam |
| VDAC (#4661) | 1:1000 | Cell Signaling |
| Grp78 (#3177) | 1:1000 | Cell Signaling |
| Ndufs1 (#70264) | 1:1000 | Cell Signaling |
| Sdha (#11998) | 1:1000 | Cell Signaling |
| Uqcrc2 (#99258) | 1:1000 | Cell Signaling |
| Cox IV (#4850) | 1:1000 | Cell Signaling |
| GAPDH (NB300-327) | 1:10000 | Novus Biological |

### Analysis of SOMs specificity for GDF15

The capability of the SOMs to bind and disrupt GDF15 dimeric form was first tested through a western blotting analysis. Briefly, 50ng of purified human recombinant GDF15 (Sigma) were incubated with 200µM of the SOMs or vehicle (DMSO) for 30min at 4°C, in slow agitation. Then, samples were prepared for SDS-PAGE and western blotting was performed, as indicated in the previous section. GDF15 antibody was used to visualize the monomeric and dimeric form of GDF15. Interference with GFRAL receptor was tested by using the GDF15:GFRAL [Biotinylated] Inhibitor Screening Chemiluminescent Assay Kit (BPS Bioscience). SOMs were tested at 200µM, following manufacturer’s instructions, and chemiluminescence was read using Synergy^TM^ fluorometer (Bio-Tek Instruments, Winooski, Vermont, USA)

### Glucose consumption and ROS production analyses

Glucose consumption and ROS production were analyzed in OV90 cells, using commercially available kits. Specifically, Glucose-Glo™ Assay was used for glucose measurement in the culture media and ROS-Glo™ H_2_O_2_ Assay for ROS measurement. Both kits were purchased from Promega.

Briefly, for glucose measurement, 8000 cells/well were seeded in a 96-well plate. After 72h of treatment with 100μM of the SOMs, an aliquot of the culture medium was taken and diluted to 1:300 in PBS. Glucose concentration was then quantified. In particular, the assay couples glucose oxidation and NADH production with a bioluminescent NADH detection system. Glucose dehydrogenase uses glucose and NAD+ to produce NADH. In the presence of NADH a pro-luciferin Reductase Substrate is converted by Reductase to luciferin that is then used by Ultra-Glo™ Recombinant Luciferase to produce light.

For ROS production measurement, 8000 cells/well were seeded in a 96-well plate. Cells were treated with 100μM of the SOMs for 72h. The kit uses a substrate that reacts with H_2_O_2_ to create a luciferin precursor. This precursor is then converted to luciferin that is then used by Ultra-Glo™ Recombinant Luciferase to produce light. Light signal is proportional to the amount of H_2_O_2_ in the sample.

All treatments were performed in triplicate, in two independent experiments pooled together, and chemiluminescence was assessed using the GloMax multiplate reader (Promega).

### Statistical analyses

Results are shown as mean ± standard deviation (SD). The Shapiro-Wilk normality test was performed to check whether the data were normally distributed. Then, Student’s t test was used to analyze the mean values after each treatment with respect to the controls. SPSS 23.0 software for Windows (SPSS Inc.; Chicago, IL, USA) was used for analyses. P values <0.05 were considered statistically significant

## Results

### *In silico* screening allows the identification of GDF15-interacting SOMs

To identify novel SOMs capable of interacting with GDF15 a structure-based virtual screening workflow was implemented. The approach combined the consideration of distinct oligomeric states, and systematic molecular docking of a curated compound library. Since GDF15 can exist in both monomeric and dimeric forms, both structural states were included in the screening pipeline. This dual approach allowed exploration of distinct binding site topologies potentially involved in ligand recognition, improving the likelihood of identifying modulators that could stabilize or disrupt specific conformations of the protein. Each form was independently subjected to docking grid generation around the putative interaction surfaces. A large library of commercially available SOMs was then docked into both structural states, SOMs were ranked according to docking scores, and the most promising candidates were selected based on both predicted binding affinity and interaction profile diversity, ensuring prioritization of chemically and functionally diverse scaffolds for subsequent *in vitro* and *in silico* analyses. Finally, sample availability from compound provider was also verified, leading to a final list of 6 SOMs, 3 for each of the different state of the target considered in the study, that were purchased and experimentally tested. List of selected SOMs is reported in **Table 2**.

**Table 2:**
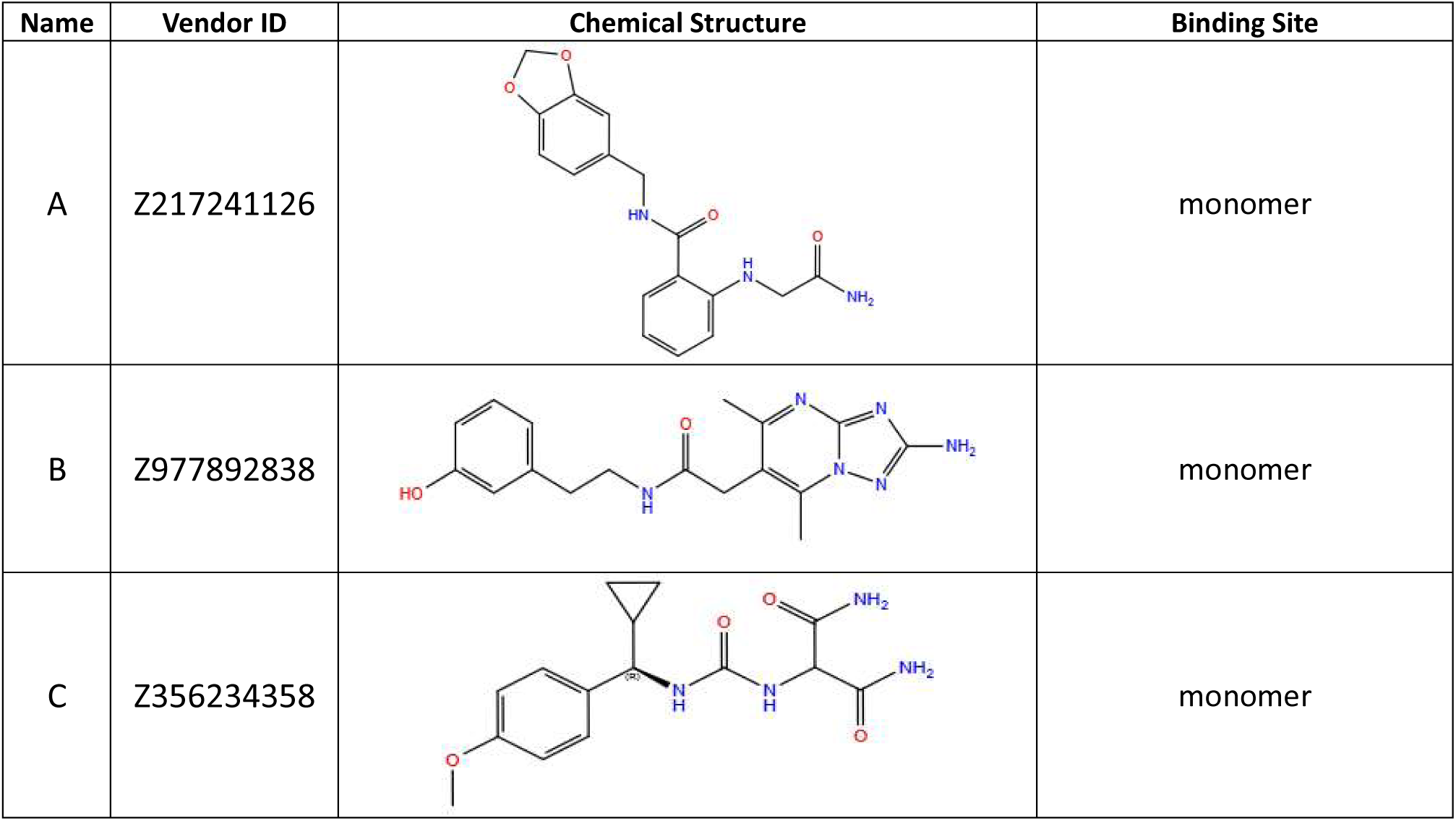
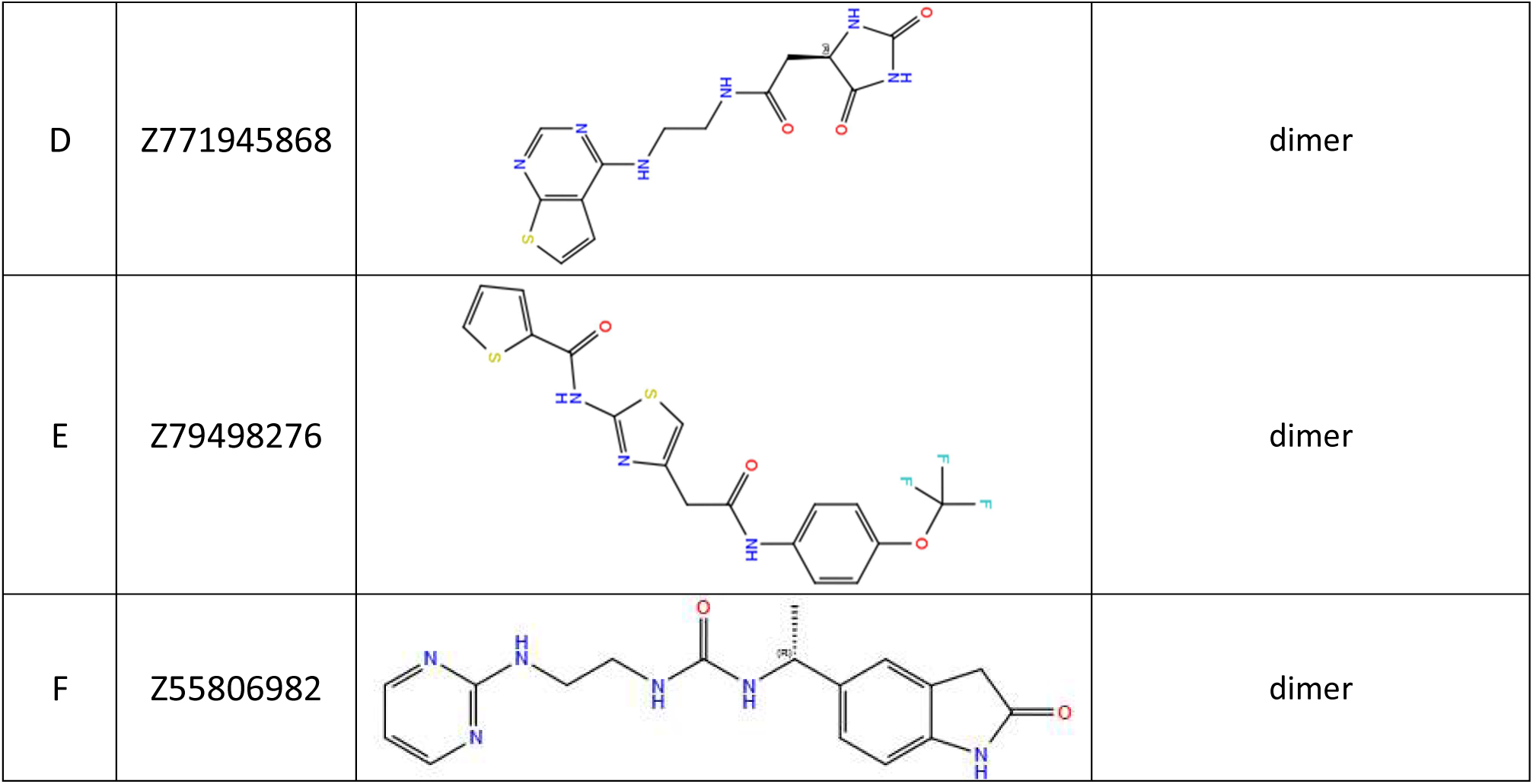
List of selected SOMs and their chemical structure.

| Name | Vendor ID | Chemical Structure | Binding Site |
| --- | --- | --- | --- |
| A | Z217241126 |  | monomer |
| B | Z977892838 |  | monomer |
| C | Z356234358 |  | monomer |
| D | Z771945868 |  | dimer |
| E | Z79498276 |  | dimer |
| F | Z55806982 |  | dimer |

### All tested SOMs reduce GDF15 dimer formation while SOM D and E also reduce GDF15-GFRAL binding

First, we tested the ability of the SOMs to reduce the abundance of the dimeric form of GDF15, considered the biologically most active form of this protein. After incubating 50ng of human recombinant GDF15 with the SOMs (200µM), we performed a western blotting analysis to evaluate the relative abundance of the dimeric and monomeric forms of GDF15. All tested SOMs reduced the abundance of the dimeric species compared to the DMSO treated control; this result indicates that the compounds could destabilize or prevent the formation of the inter-chain disulphide linkage required for dimerization. Although the magnitude of the effect varied among molecules, the overall trend was consistent, suggesting that each SOM interacts with regions critical for dimer assembly or stability (**Fig. 1A)**.

**Figure 1.**
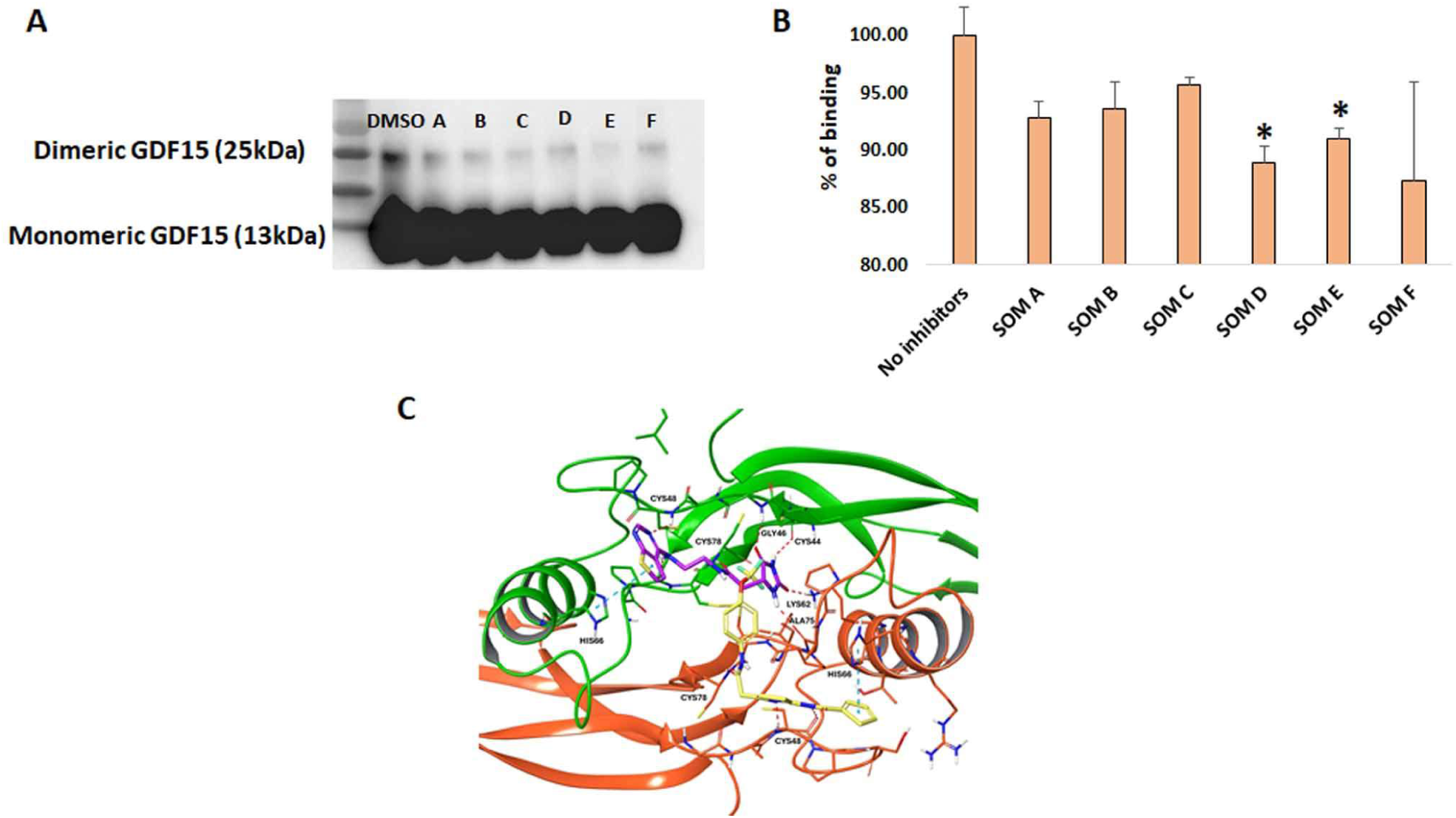
Analysis of SOMs specificity for GDF15. (**A**) Western blotting analysis of human recombinant GDF15 (50ng), incubated with each SOM (200μM) or the solvent (DMSO), for 30 minutes. Both dimeric and monomeric forms of GDF15 are visible. (**B**) Chemiluminescence assay, testing the ability of the SOMs to disrupt GDF15-GFRAL binding. Percentage of binding compared to control is showed in the graph; each condition was tested in triplicate. Data are expressed as mean ± SD. Student’s t test was applied. * p< 0.05. (**C**) Predicted binding mode of compounds D (purple) and E (yellow). The two GDF15 monomers are colored respectively green and orange, interacting residues are drawn as thick tubes and labeled. Hydrogen bonds are represented as red dotted lines, pi-pi interactions as blue dotted lines.

We then sought to analyze the binding of GDF15 to GFRAL under SOMs treatment. To do so, we used a commercial chemiluminescent assay, which uses streptavidin-HRP to detect biotin-labeled GFRAL, that binds to GDF15-coated wells. A moderate but statistically significant decrease in receptor binding was observed for SOMs D and E (**Fig. 1B)**. These compounds produced the most pronounced inhibition, consistent with a potential ability to interfere with the receptor-recognition surface of GDF15. The absence of a marked effect for the remaining SOMs implies either binding outside the receptor interface or insufficient affinity to perturb GDF15–GFRAL association under the assay conditions. SOMs D and E were originally identified through virtual screening against the dimeric form of GDF15, and their ability to impair GDF15–GFRAL interaction can be explained by considering the location of the predicted binding site. **Figure 1C** illustrates the predicted binding mode of molecules D and E within the interfacial cavity of the GDF15 dimer; both ligands occupy overlapping regions at the dimer interface, forming a dense network of hydrogen bonds and hydrophobic contacts with residues critical for structural stabilization. Interestingly, due to the symmetry of the active site cavity, the two ligands are predicted to form key interaction with same residues but in different GDF15 subunit, such as the hydrogen bond interactions with Cys48 and Cys78, and the pi-pi interactions with His66. The positioning of both molecules along the inter-monomer cleft suggests that they could act by perturbing the local geometry of this interface, thereby potentially modulating the relative orientation of the two subunits.

### Effect of SOMs on viability and proliferation of OV90 cells and DFs

To evaluate the potential effects of SOMs *in vitro*, we used two different cell models, a normal primary cell line and a tumor cell line, since it is already reported in the literature that tumor cells express higher levels of GDF15 with respect to normal cells (**Li et al, 2018**; **Hüllwegen et al, 2025).** As expected, OV90, a human ovarian cancer cell, and human primary dermal fibroblasts (DFs), are characterized, among others, by a very different basal expression level of GDF15. As showed in **Suppl. Fig. 1**, OV90 showed a much higher expression of *Gdf15* than DFs.

We performed in both models a proliferation and viability test in a 72h-time course, assessing viability and total number of cells treated with each SOM at 100µM at 24, 48 and 72h. At 24 and 48h no evident effects were observed in OV90 cells, except for SOM E, which significantly reduced viability and proliferation, compared to control (**Fig. 2 A, B**). The effect of SOM E was even more evident at 72h; at this same time point, also SOMs C, D and F reduced the total number of cells, compared to controls, although SOM C not significantly (**Fig. 2 B)**. On DFs, after a 72h-treatment, no significant effects were observed except for SOM E that tended to reduce viability (data not shown). Considering these preliminary results, we decided to focus our subsequent analyses on SOMs C, D and F. SOM E was excluded due to its toxicity on cell cultures and separately tested in different experiments (*see below*).

**Figure 2.**
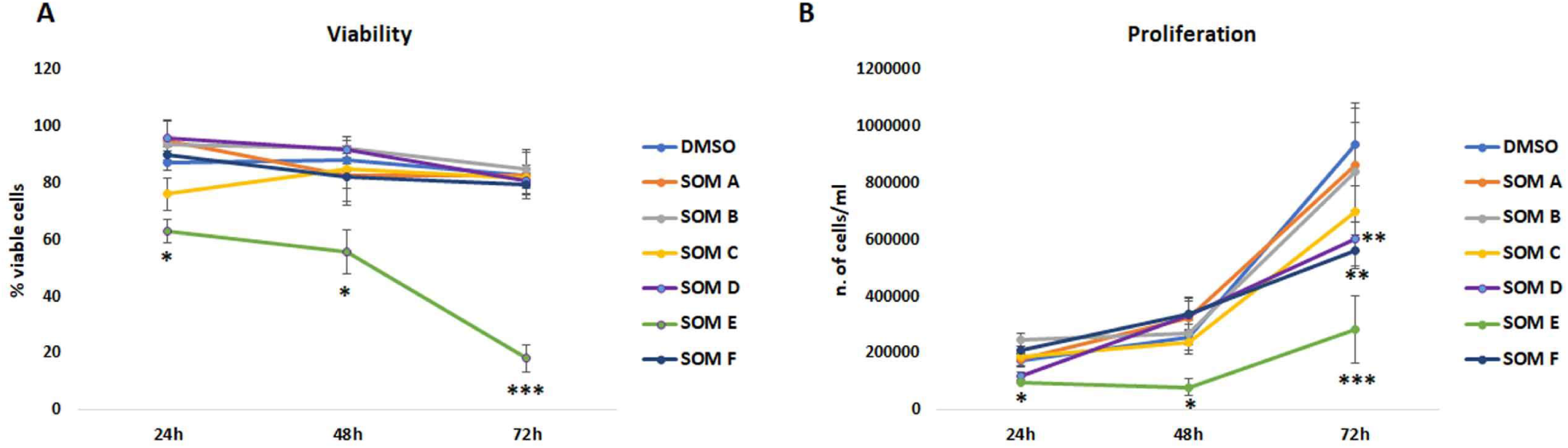
Viability and proliferation analysis of OV90 cells after small organic molecules (SOMs) treatment. (**A**) Viability assessed at 24, 48, 72h, by staining cells with Trypan blue and using an automated cell counter. (**B**) Proliferation assessed by counting the total number of cells at each time point, using an automated cell counter. Each condition was performed in quadruplicate. Data are expressed as mean ± SD. Student’s t test was applied to compare each SOM treatment with respect to DMSO at each time point. * p< 0.05; ** p< 0.01; *** p< 0.001

### Modulation of gene expression is mostly observed in OV90 cells upon SOM D treatment

In order to investigate possible effects of SOMs C, D and F on gene expression, we performed quantitative real time RT-PCR analyses on RNA extracted from both cell lines treated with 100µM of the selected SOMs for 72h. First, we evaluated the expression of genes related to cell cycle and proliferation. *p21* expression was significantly higher in OV90 treated with SOM C and D, compared to DMSO treated control cells **(Fig. 3 A)**. In DFs, no significant differences were observed **(Fig. 3 B).** *Ki67* expression was reduced by SOM D treatment in both OV90 **(Fig. 3 C)** and DFs (**Fig. 3 D).**

**Figure 3.**
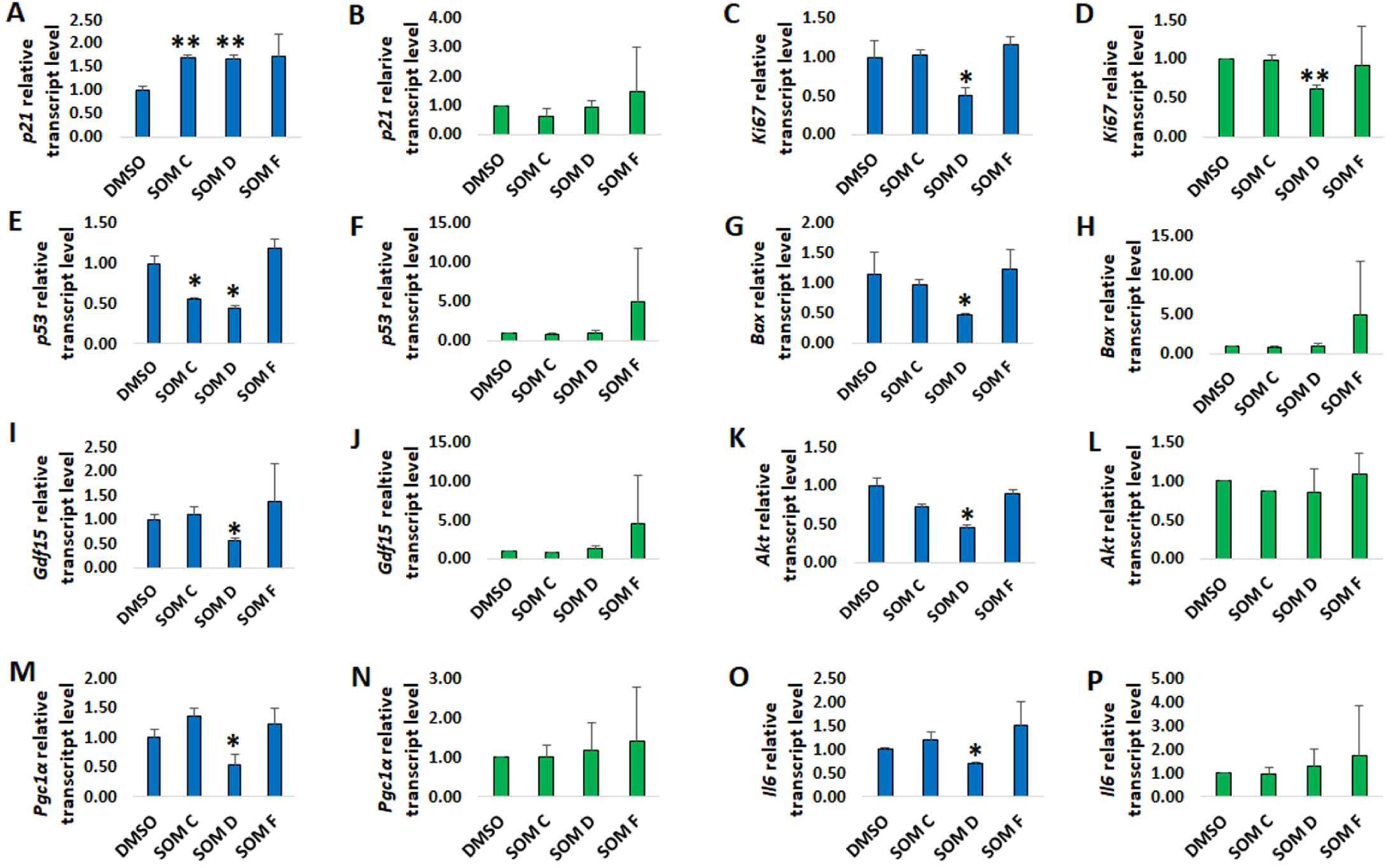
Real time RT-PCR analysis in OV90 (blue columns) and primary dermal fibroblasts (green columns), after 72h-SOMs treatment. Analysis of (**A-B**) *p21,* (**C-D**) *Ki67,* (**E-F**) *Bax,* (**G-H**) *p53,* (**I-J**) *Gdf15,* (**K-L**) *Akt*, (**M-N)** *Pgc1a* and (**O-P**) *Il6* expression in 3 OV90 replicates and 2 fibroblasts lines. Data are expressed as mean ± SD. Student’s t test was applied to compare each small organic molecule (SOM C, D, F) treatment with respect to DMSO. * p< 0.05; ** p< 0.01

We also analyzed the expression of *p53* and *Bax*, two genes related to apoptosis. Moreover, p53 is also a GDF15 transcription factor (**Osada et al, 2007; Conte et al, 2022).** *p53* expression was reduced in OV90 treated with SOM C and SOM D (**Fig. 3 E).** In DFs, SOM F determined a strong, but not significant, increase in *p53* expression (**Fig. 3 F).** As regards *Bax* expression, SOM D reduced its expression in OV90 (**Fig. 3 G).** In DFs, SOM F determined a similar trend to that observed for *p53* (**Fig. 3 H)**.

We then evaluated the expression level of *Gdf15* itself and *Akt*, a molecule involved in GDF15 signaling pathway (**Takahashi, 2022**). Surprisingly, in OV90 cells, SOM D determined a reduced expression of both *Gdf15* and *Akt* (**Fig. 3 I, K)**, while in DFs no significant variations were observed **(Fig. 3 J, L).** It is not known whether GDF15 can control the expression of Akt, therefore, in order to verify whether the effect of the SOM D was really due to GDF15 downregulation, we performed the same analysis upon a RNAi on OV90 cells, using a siRNA targeting GDF15. About 70% of *Gdf15* expression was effectively reduced by siRNA treatment (**Suppl. Fig 2 A)** and, similarly to what was observed with SOM D, GDF15 KD caused a downregulation of *Akt* expression, as well as of *Ki67* **(Suppl. Fig. 2 B, C).**

GDF15 is reported to be strongly associated with mitochondrial stress, with reported mitochondria-protective effects. Moreover, both pro- and anti-inflammatory effects have been associated to GDF15 (**Conte et al, 2022; Breit et al, 2021; Bonaterra et al, 2012; Luan et al, 2019**). Therefore, we evaluated the expression of *Pgc1α,* a gene involved in mitochondrial biogenesis and *Il6*, an inflammation-associated cytokine. *Pgc1α* expression was significantly reduced by SOM D treatment in OV90 (**Fig. 3 M)**, while no significant variations were observed in DFs (**Fig. 3 N**). As regards *Il6,* its expression was reduced in OV90 after SOM D treatment **(Fig. 3 O).** No significant differences were found in DFs **(Fig. 3 P).**

Finally, in order to evaluate the possibility to use SOM E at non-toxic doses, we performed a dose-response curve for this SOM. Viability and proliferation were not affected by doses 1 and 10 μM (data not shown), however, when testing the expression of 7 out of 8 of the above-mentioned genes (*p21, Ki67, p53*, *Bax, Gdf15, Akt and Il6*), the results indicated that for these genes, lower doses had no significant effect, while the 100 μM dose caused a different effect (or no effect) with respect to other SOMs (a much higher increase of *p21* and an increase in *Gdf15, p53*, *Bax*, *Akt* and *Il6* expression, instead of a decrease) (**Suppl. Fig. 3**). This suggests that the mechanism of action of SOM E is likely different from that of other SOMs, in particular SOM D.

Overall, these data suggest that SOM D may exert biological effects consistent with GDF15 inhibition. Interestingly, these effects are evident in OV90, a cancer cell line with a high proliferation rate and a high basal level of *Gdf15* expression (see **Suppl fig. 1)** but not in normal primary cell lines such as DF.

### SOM D determines an alteration of mitochondrial proteins expression

We continued the characterization of the biological effects of the SOMs, specifically testing whether SOMs treatments induced stress at endoplasmic reticulum or mitochondrial level, thus impinging upon GDF15 processing. To this end, we performed a western blotting analysis on OV90 cells, treated for 72h with SOMs C, D, and F. First, we analyzed the protein expression of GDF15, as a downstream of the mitochondrial Unfolded Protein Response. Both its precursor (pro-GDF15) and the mature form (m-GDF15) were detectable **(Fig. 4 A)**. We calculated the m-GDF15/pro-GDF15 ratio in order to have an indication of GDF15 processing. This ratio was significantly higher after treatment with SOM C (**Fig 4 B).** Then, we analyzed the protein expression of Grp78, considered a marker of endoplasmic reticulum stress, and of VDAC, used as a marker of mitochondrial mass. No significant variation of Grp78 were observed, after treatment with the SOMs **(Fig. 4 A-C)**. Interestingly, VDAC amount upon GAPDH normalization was statistically higher in cells treated with SOM D, compared to DMSO treated cells (**Fig. 4 A-D)**.

**Figure 4.**
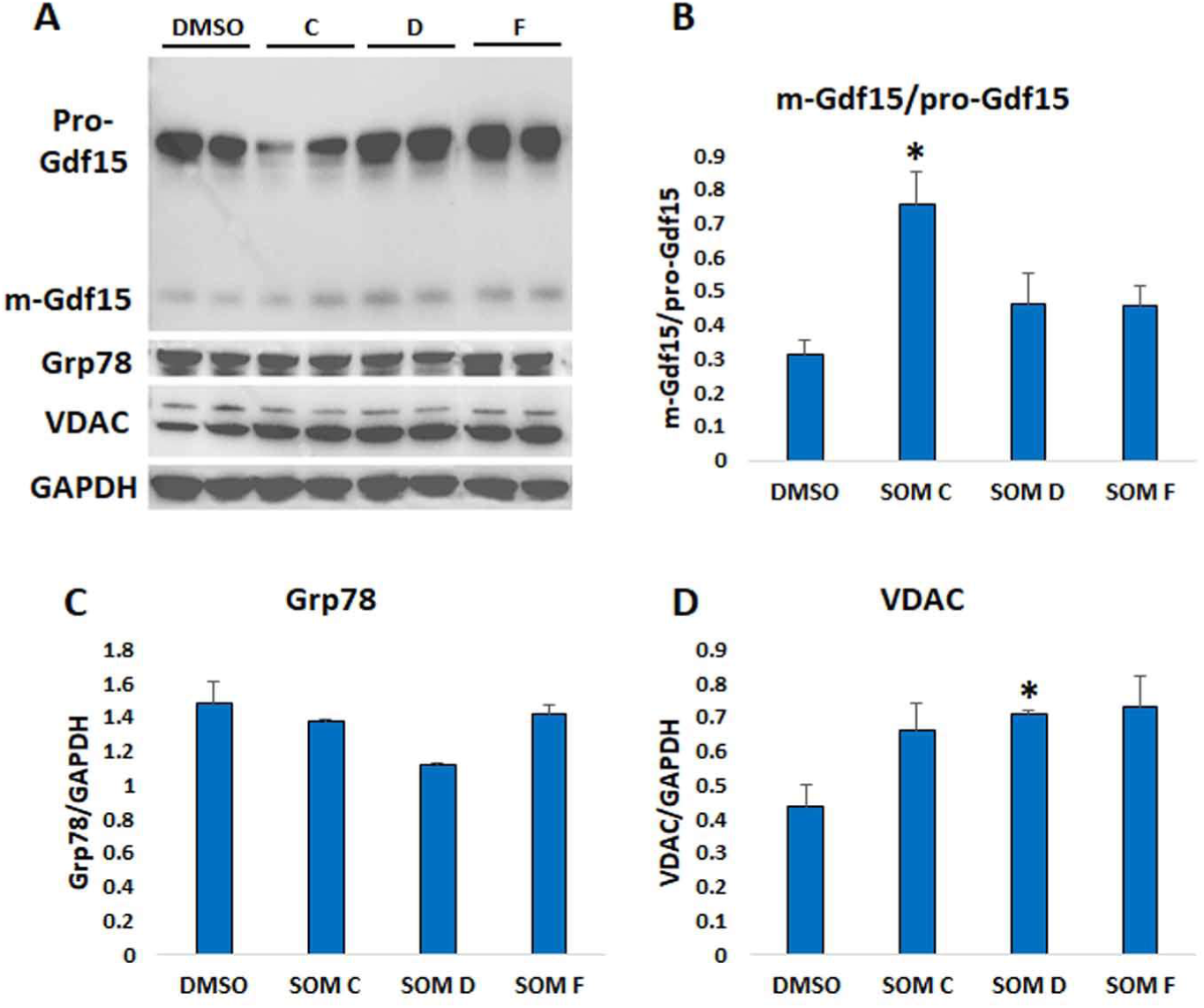
Western blotting analysis of Gdf15, Grp78, VDAC and GAPDH in OV90, after 72h-treatment with SOM C, D and F. (**A**) Representative immunoblotting image of Gdf15 (m-Gdf15: mature form; pro-Gdf15: precursor form), Grp78, VDAC and GAPDH. (**B**) m-Gdf15/pro-Gdf15 ratio, based on m-Gdf15 and pro-Gdf15 band intensity. (**C**) Grp78 relative protein expression. (**D**) VDAC relative protein expression. Data are expressed as mean ± SD. Student’s t test was applied to compare each SOM treatment with respect to DMSO. * p< 0.05. The quantification was performed using Fiji software and normalized to GAPDH expression.

Finally, we tested the protein expression of representative subunits of the mitochondrial complexes. SOM D reduced the expression of Sdha and Cox IV, whereas SOM F reduced the expression of Sdha. None of the SOMs caused variations of Ndufs1 and Uqcrc2 expression (**Fig 5 A-E).** These data suggest that SOM D in particular may cause an impairment of mitochondrial complexes assembly, in agreement with the observed modulation of *Pgc1α* expression.

**Figure 5.**
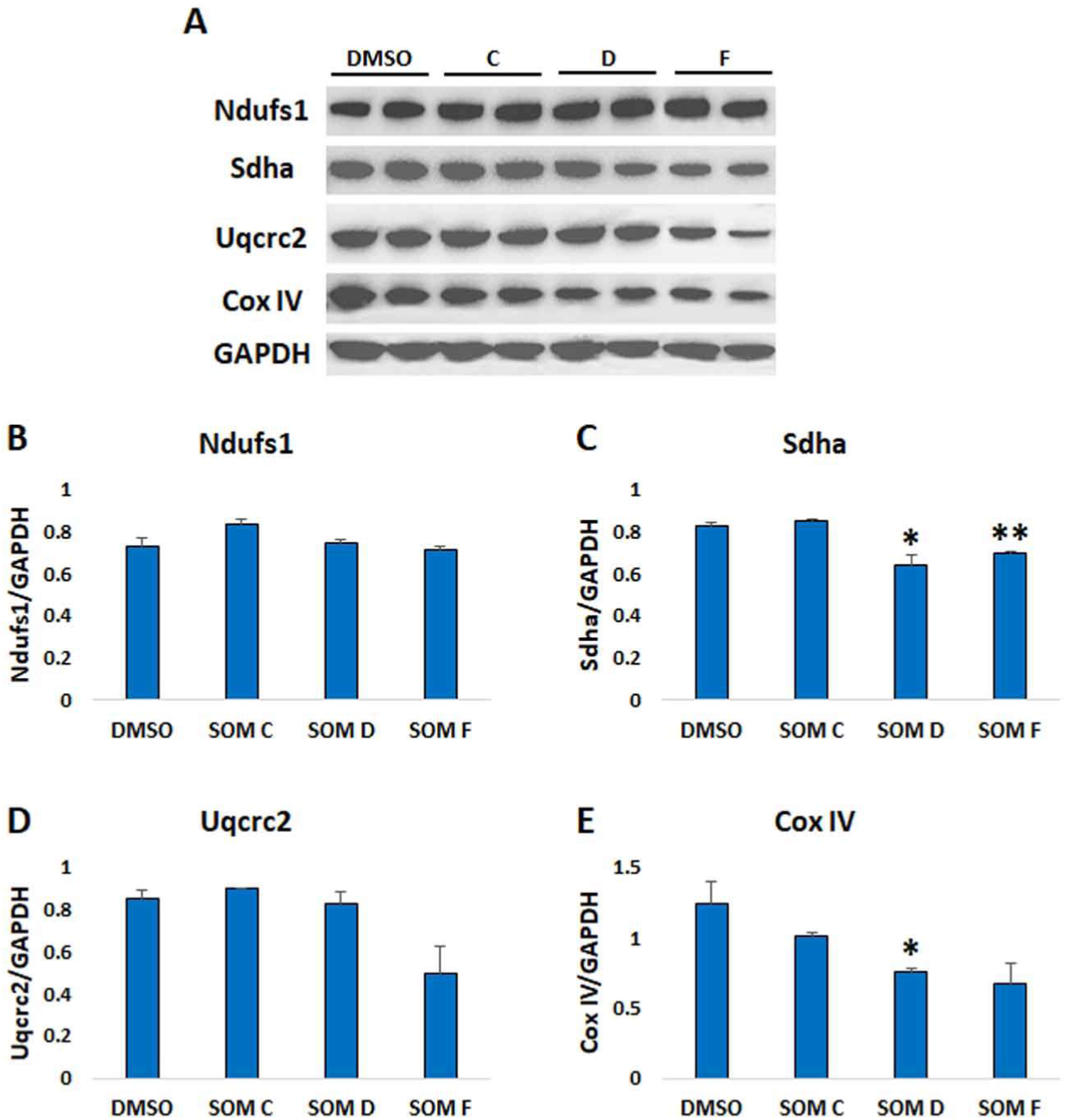
Western blotting analysis of Ndufs1, Sdha, Uqcrc2 and Cox IV in OV90 after 72h-treatment with small organic molecule (SOM C, D and F). (**A**) Representative immunoblotting image of Ndufs1, Sdha, Uqcrc2, Cox IV and GAPDH. Relative protein expression of (**B**) Ndufs1, (**C**) Sdha, (**D**) Uqcrc2 and (**E**) Cox IV. Data are expressed as mean ± SD. Student’s t test was applied to compare each SOM treatment with respect to DMSO. * p< 0.05, **p<0.01. The quantification was performed using Fiji software and normalized to GAPDH expression.

### SOMs C and F increase ROS production in OV90 cells

Considering that SOMs C, D and F affected cell proliferation, gene and protein expression in OV90 cell line, we sought to verify if these molecules also affected glucose consumption and ROS production. To do so, we used two commercially available kits to assess, via a chemiluminescence reaction, glucose concentration and H_2_O_2_ production in OV90, after a 72h-treatment with the above-mentioned SOMs. Results were normalized over the total number of cells. SOMs C and D tended to increase glucose consumption, although not significantly (**Fig. 6 A).** As far as ROS production, a small but significant increase in H_2_O_2_ production was observed in cells treated with SOMs C and F (**Fig. 6 B)**. Overall, these data indicate that SOMs C and F but not D could potentially affect the redox cellular balance.

**Figure 6.**
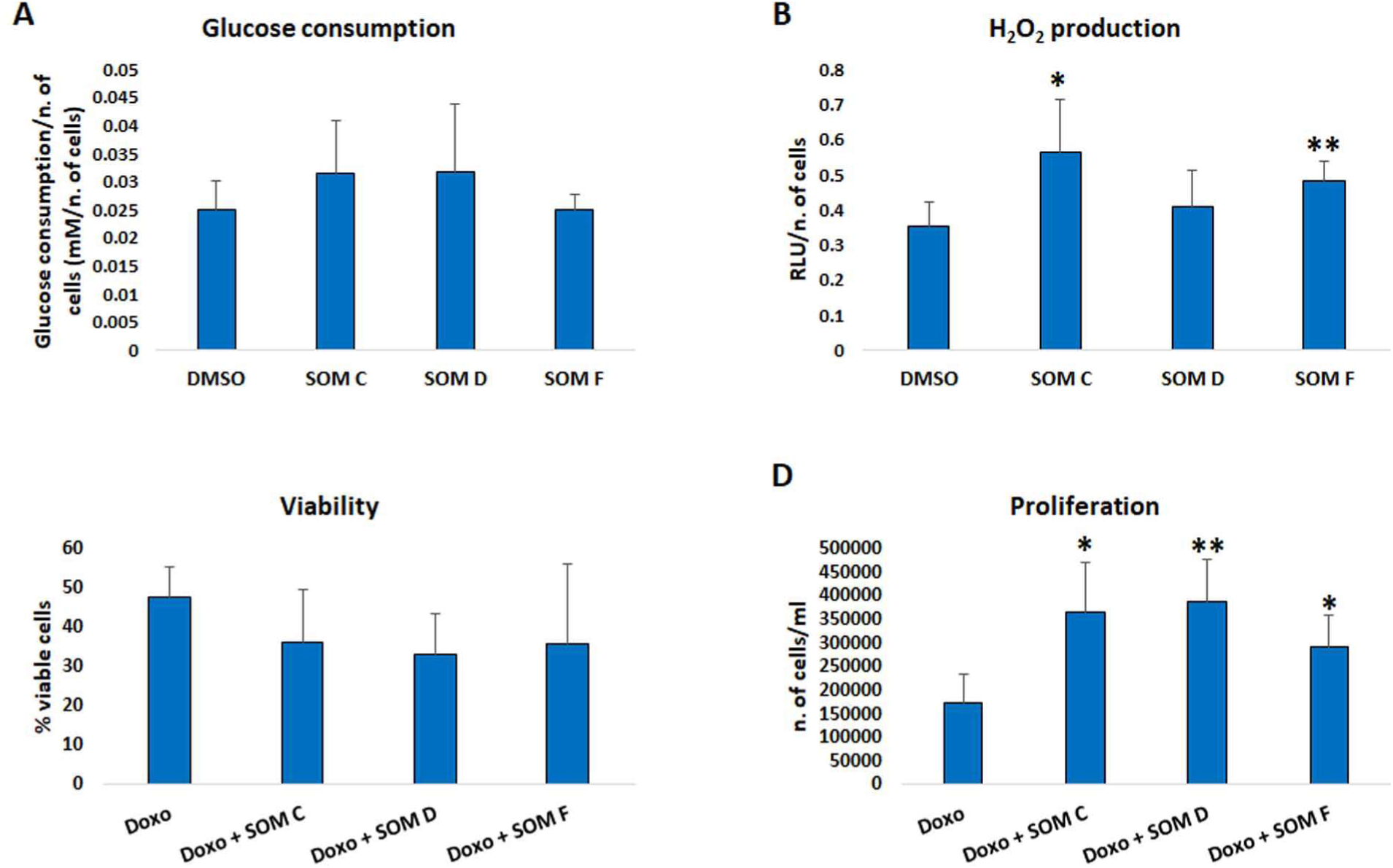
Functional assays and viability and proliferation analysis of OV90 cells. (**A**) Glucose consumption and (**B**) ROS (H_2_O_2_) production chemiluminescence analysis. Glucose consumption is expressed as the difference between the initial glucose concentration in the culture medium, in mM, and the concentration at the end of the treatment. This value is then normalized on the total number of cells. H_2_O_2_ production is expressed as relative light unit (RLU) per cell. Each condition in both assays was performed twice in triplicate. (**C**) Viability and (**D**) proliferation after 24h pre-incubation with 1μM doxo followed by 72h SOMs treatment, assessed by staining cells with Trypan blue and using an automated cell counter. Each condition was performed in quadruplicate. Data are expressed as mean ± SD. Student’s t test was applied. * p< 0.05; ** p< 0.01.

### SOMs C, D and F reduce doxorubicin effect on OV90 proliferation

We then tested the effect of SOMs C, D and F on OV90 cells when treated with doxorubicin, a widely used chemotherapeutic drug. We performed a 24h pre-incubation with 1μM doxorubicin, then the medium was replaced and 100μM of each SOM was added for further 72h. Co-treatment with doxorubicin + SOM C, SOM D or SOM F tended to reduce cell viability compared to doxorubicin alone, although not significantly (**Fig 6 C**). Unexpectedly, the total number of cells was significantly increased after co-treatment with doxorubicin + SOM C, SOM D or SOM F, compared to doxorubicin alone **(Fig 6 D**). This suggests that these molecules may inhibit the cell cycle blockade caused by doxorubicin.

## Discussion

Elevated plasma levels of GDF15 have been associated with several pathologies, including T2D, cardiovascular, renal, and neurodegenerative diseases, as well as sarcopenia and frailty (**Conte et al., 2022; Conte et al., 2025**). Therefore, the possibility to modulate GDF15 activity seems a potential treatment option for a number of disorders. Accordingly, GDF15 role in cancer cachexia is particularly well documented, and promising data on the use of moAbs against GDF15 in clinical trials with patients with advanced cancer have been recently reported (**Groarke et al., 2024; Melero et al., 2025**). In this study, we aimed to explore the possibility of applying an *in silico* screening methodology to discover commercially available SOMs potentially able to bind GDF15 in monomeric or dimeric form and modulate its biological effects, thus paving the way for a novel strategy for GDF15 inhibition.

The identified SOMs appear to reduce GDF15 dimer formation, and two of them—molecules D and E—also significantly decrease its binding to GFRAL. Both compounds were discovered through virtual screening performed on the dimeric structure of GDF15, and their behavior can be explained by the location of their predicted binding site. Docking indicated that the molecules bind within a cavity at the interface of the two monomers, a region likely important for maintaining the dimer’s overall shape and the conformation of the receptor-binding surface. Binding of SOMs D and E in this pocket may cause local conformational adjustments that spread to the external parts of the dimer, slightly altering the geometry of the GFRAL-binding region and reducing receptor recognition. Because the screening was performed on a pre-assembled dimer, and even the less effective molecules (*e.g.*, SOM A and B) decreased dimer abundance, D and E probably do not inhibit GDF15 by preventing dimerization. Instead, they seem to act as allosteric modulators: by occupying the inter-monomer cavity, they stabilize a dimeric form that remains structurally intact but is less compatible with GFRAL binding. This model may explain the reduced GDF15–GFRAL interaction, suggesting that SOMs D and E may function as allosteric antagonists. Further experiments will be required to confirm this hypothesis.

No effects on cell viability were observed, except for SOM E that resulted acutely toxic, likely through mechanisms that go beyond GDF15 inhibition. For this reason, it was excluded from further testing. Proliferation was reduced in OV90 upon treatment with SOM D and F after 72h of treatment, while in DFs no effect was observed. This difference in the susceptibility to SOMs may be due to i. the different levels of GDF15 expressed by the two experimental models considered, and therefore to a different dependence on GDF15; ii. their different proliferation’ rate. Further studies on a longer exposure time will be needed to exclude the presence of long-term effects on normal cells like DFs.

As far as gene expression, upon treatment with SOM D, OV90 display an increased expression of *p21* and a decrease in *Ki67*, consistently with the data on proliferation, a decreased expression of *Akt* (a molecule part of GDF15 signaling pathway), *p53*, *Bax* and *Gdf15* itself. Mitochondria appear impaired (mass is increased, and biogenesis is likely altered as *Pgc1*α, Sdha, and Cox IV expression are decreased); however, H_2_O_2_ production and glucose consumption are unaffected. At variance, SOM C and F caused an apparently smaller effect on mitochondrial subunit expression but a significant increase in H_2_O_2_ production. Since all these three SOMs cause a decrease in proliferation (although just as a trend for SOM C), it is likely that this is linked to an effect on mitochondrial function for SOM C and F but not SOM D, which appears to be linked to increased p21 expression and decreased Ki67. Previous report has linked GDF15 to cell senescence (**Chiariello, Rossetti et al., 2024**). Accordingly, SOM D appears to affect cell cycle much more than mitochondrial function. Interestingly, the effects are seen primarily in OV90 cancer cells that express high levels of GDF15; almost nothing is observed in normal cells such as DFs.

While we have no indication whether these SOMs (especially D) have an anticachectic effect, for which data from *in vivo* models would be needed, we do have an interesting indication on which cells, normal or cancer ones, are more affected by inhibition of GDF15 signaling. In particular, according to our data, normal cells do not appear to be significantly affected by treatment with SOMs at these doses and times, unlike tumor cells, which increase the expression of senescence markers such as p21 and decrease the expression of GDF15 itself. This suggests that the use of SOMs against GDF15 at the cellular level should be non-toxic for normal cells, also in agreement with what was observed in clinical trials with anti-GDF15 moAbs, where no increase in side effects was recorded in the treated patient group compared to the placebo group **(Groarke et al., 2024**).

The beneficial effect of GDF15 signalling inhibition on cancer progression and cachexia in vivo is pretty well established (**Suriben et al., 2020; Groarke et al., 2024; Melero et al., 2025; Shi et al, 2025**). The effects of GDF15 signaling inhibition on single cells are much less known, with controversial findings. Our results suggest that GDF15 likely plays a complex role in cell cycle control that deserves further investigation. Indeed, our data indicate that in the absence of other stresses, SOM C, D and F induce a decrease in proliferation, while in combination with a DNA-damaging agent such as doxorubicin, they induce a smaller decrease than doxorubicin alone. This is in agreement with our previous study where the silencing of GDF15 in control or stressed cells produced an induction or a decrease of cell senescence, respectively (**Chiariello, Rossetti et al., 2024**). This may suggest that GDF15 represents a sort of checkpoint for cell cycle progression. If GDF15 is inhibited, the cell, even if damaged, continues to proliferate. This effect, if confirmed by further studies, would suggest caution on the use of GDF15 inhibitors during anticancer therapy with DNA-damaging agents.

Overall, SOM D (corresponding to vendor’s code Z771945868) seems to be a promising candidate molecule for the inhibition of GDF15 activity. Other SOMs, have only partial effects or display a significant toxicity suggesting a different mechanism of action other than GDF15 signaling inhibition.

## AUTHOR CONTRIBUTIONS

AC performed and designed experiments, maintained cell cultures, performed statistical analyses and wrote the manuscript. LT maintained cell cultures and performed Real Time PCR analyses. MS and ML critically discussed the results. ADR, LL and MDP performed *in silico* analysis and provided SOMs. SS and MC carried out the design of the study, critically analyzed the results, and wrote the manuscript. All authors contributed to the article and approved the submitted version.

## FUNDING INFORMATION

The work was co-funded from the Italian Ministry of University and Research (MUR) PRIN 2022, Project # 2022KS8T4N “GDF15 as a key player and a potential target to tackle ageing and age-associated diseases: an in silico, in vitro and ex vivo study” to SS and MDP.

The work was also co-funded from Next Generation EU, in the context of the National Recovery and Resilience Plan, Investment PE8 –Project Age-It: “Ageing Well in an Ageing Society” to SS. This resource was co-financed by the Next Generation EU [DM 1557 11.10.2022].

The work reported in this publication was funded by the Italian Ministry of Health, RC-2025/94 project.

The views and opinions expressed are only those of the authors and do not necessarily reflect those of the European Union or the European Commission. Neither the European Union nor the European Commission can be held responsible for them.

---

**Suppl. Figure 1.**
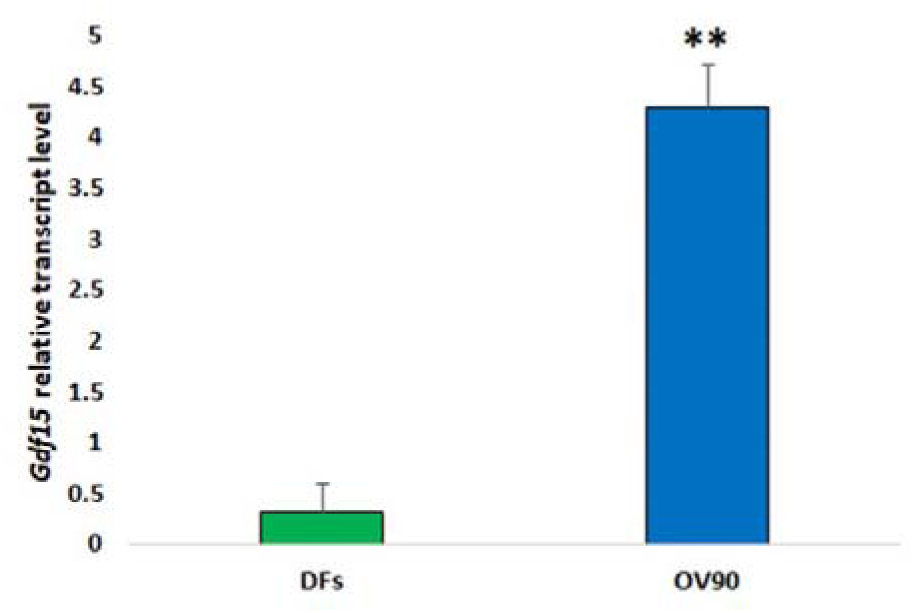
Real time RT-PCR analysis of Gdf15 basal expression in 2 lines of dermal fibroblasts (DFs) and OV90 (analysis performed in triplicate). Data are expressed as mean ± SD. Student’s t test was applied. ** p< 0.01

**Suppl. Figure 2.**
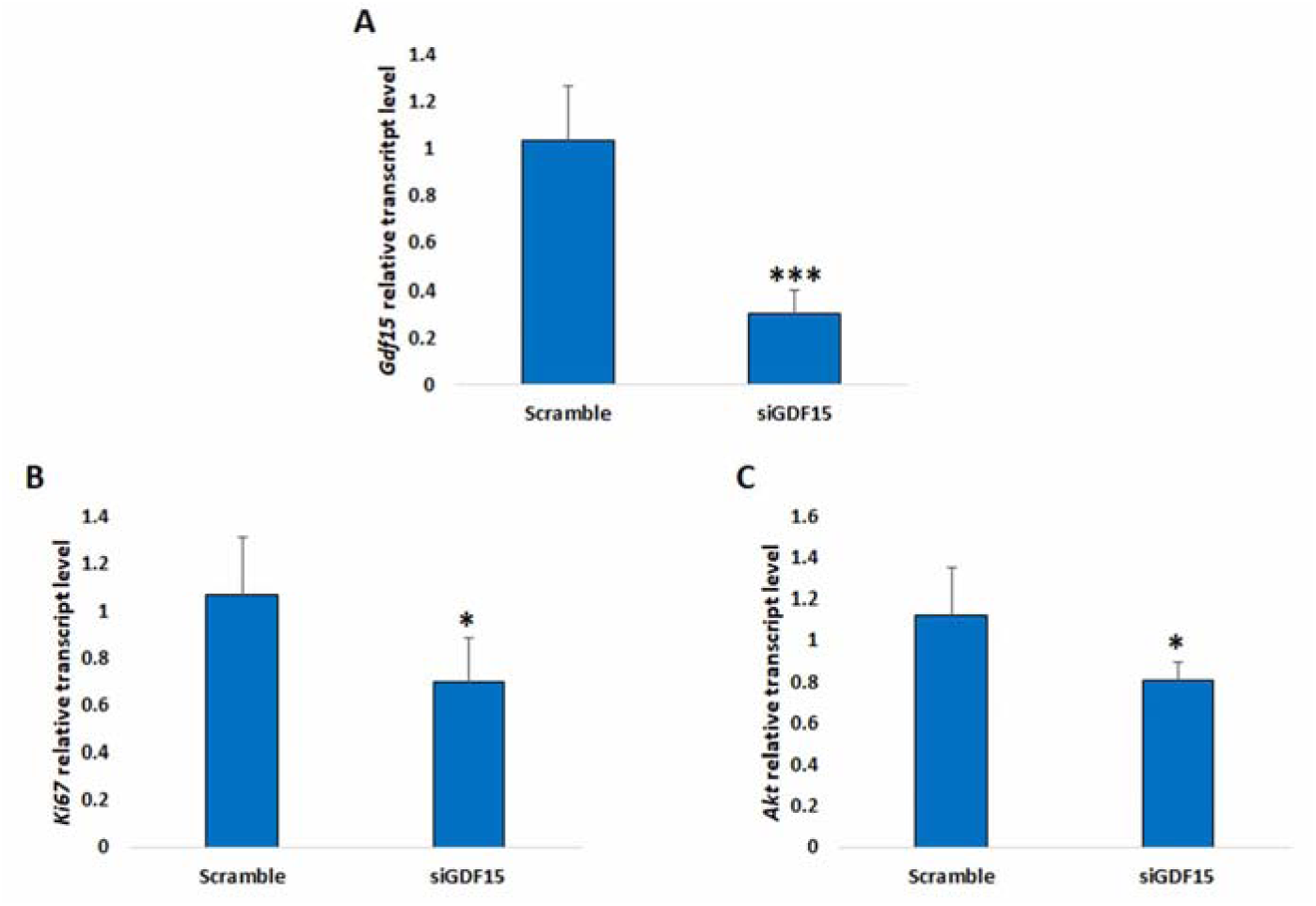
Real time RT-PCR analysis in OV90 after GDF15 knock down (siGDF15), performed in triplicate (Scramble: control siRNA). (A) Gdf15, (B) Ki67 and (C) Akt expression. Data are expressed as mean ± SD. Student’s t test was applied. *p< 0.05; *** p< 0.001

**Suppl. Figure 3.**
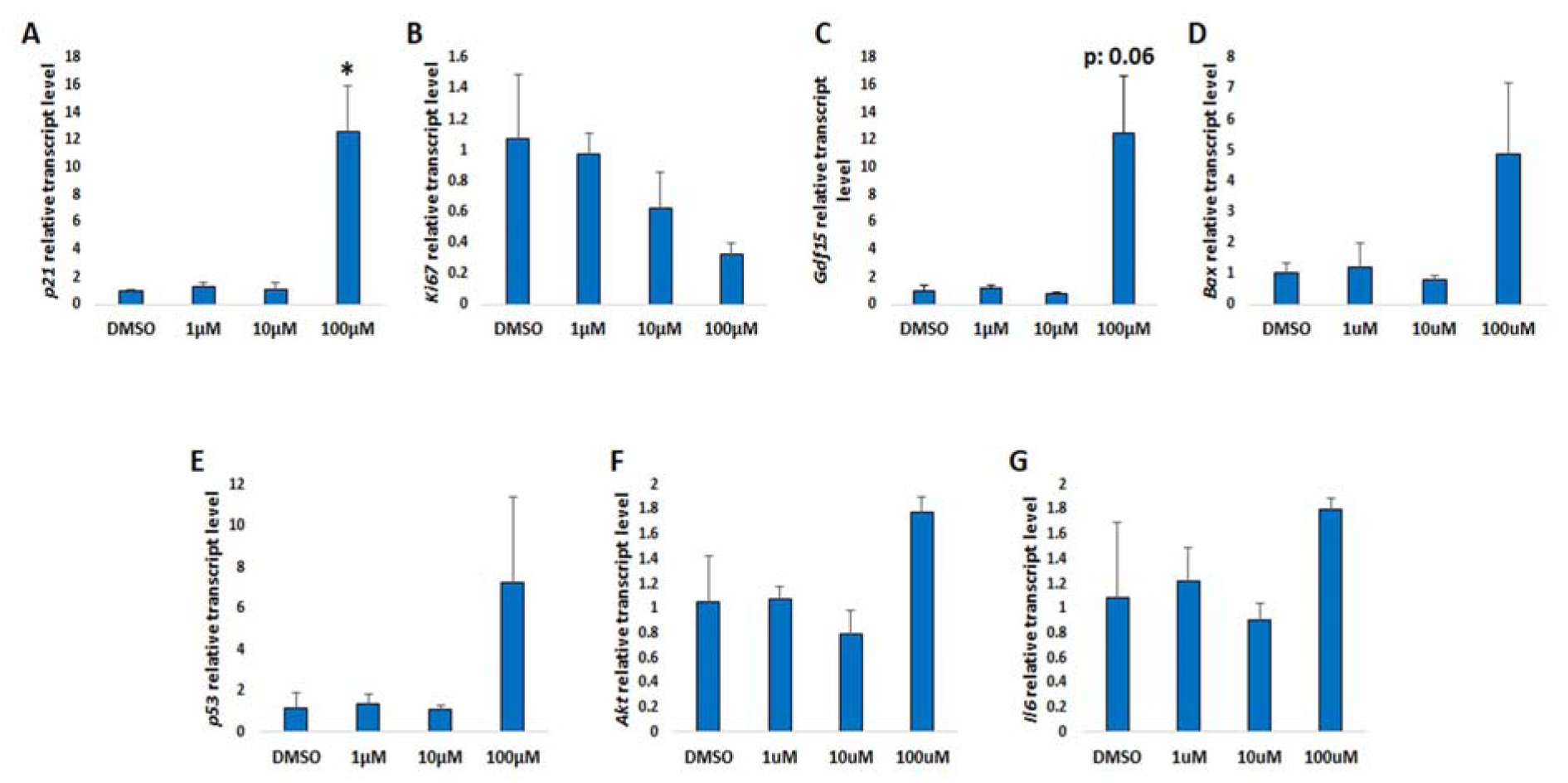
Real time RT-PCR analysis in OV90 treated for 72h with different doses of small organic molecule (SOM) E. Analysis of (A) p21, (B) Ki67, (C) Gdf15, (D) Bax, (E) p53, (F) Akt and (G) Il6 expression in OV90, performed in triplicate. Data are expressed as mean ± SD. Student’s t test was applied to compare each dose of SOM E treatment with respect to DMSO. * p< 0.05

